# Transcriptome characterization analysis and molecular profiles of obligatory diapause induction of the Chinese citrus fruit fly, *Bactrocera minax* (Diptera: Tephritidae)

**DOI:** 10.1101/672642

**Authors:** Zhixiong Zhou, Xiaolin Dong, Chuanren Li

## Abstract

The Chinese citrus fruit fly, *Bactrocera minax*, is a devastating citrus pest in China, Bhutan and India. It will enter obligatory pupal diapause in each generation at specific stage, while little is known about the course and the molecular mechanisms of diapause induction. To gain insight into possible mechanisms of obligatory pupal diapause induction, high-throughput RNA-seq data were generated from second-instar larvae (2L), third-instar larvae (3L) and pupal (P, one week after pupating). A total of 116,402 unigenes were assembled and researched against public databases, and 54,781 unigenes matched to proteins in the NCBI database using the BLAST search. Three pairwise comparisons were performed, and significantly differentially regulated transcripts were identified. Several differentially expressed genes (DEGs) expression patterns revealed that those highly or lowly expressed genes in pupal stage were predicted to be involved in diapause induction. Moreover, GO function and KEGG pathway analysis were performed on all DEGs and showed that 20-hydroxyecdysone (20E) biosynthesis, insulin signaling pathway, FoxO signaling pathway, cell cycle and metabolism pathway may be related to the obligatory diapause of the Chinese citrus fruit fly. This study provides valuable information about the Chinese citrus fruit fly transcriptome for future gene function research, and contributes to the in-depth elucidation of the molecular regulation mechanism of insect obligatory diapause induction.

## INTRODUCTION

The Chinese citrus fruit fly, *Bactrocera minax* (Enderlein) (Diptera: Tephritidae), is an important economic pest of citrus in China, Bhutan and India (Dorij *et al.* 2006; Wang and Luo 1995), and serious yield losses was caused by larval feeding (Lv *et al.* 2010; Han *et al.* 2011). This insect exhibits obligatory pupal diapause to overwinter in each generation, regardless of the prevailing environmental conditions. A number of prior studies about control methods, population dynamics, adult development have been carried out (Chen *et al.* 2012; Dong *et al.* 2014b; Dong *et al.* 2013; Gao *et al.* 2013; Wang *et al.* 2014; Zhang *et al.* 2014; Wang *et al.* 2018). And some aspect of diapause are also well established in this species, for instance, RNA sequencing (RNA-seq) was applied to investigate the transcriptome characterization differences among early diapause, late diapause and post-diapause (Dong *et al.* 2014a; Wang *et al.* 2016; Wang *et al.* 2017). However, little work has been performed to elucidate the molecular basis of diapause induction in this species.

Diapause is an alternative life history stage that allows insects to mitigate acute environmental stresses (Denlinger 2002; Koštál 2006). It is divided into three main phase: pre-diapause (including induction phase and preparation phase), diapause (including initiation, maintenance and termination) and post-diapause (Koštál 2006). Insect species enter diapause in different ontogenetic stages. Phenotypic features of diapause induction are also different among most insect species. There may be diverse transcriptional strategies for producing them. Facultative diapause occurs in response to environmental cues (including photoperiod and temperature), but obligatory diapause occurs during each generation regardless of the environmental cues it receives (Denlinger 2009). In facultative diapause insects, some studies have released the molecular basis of diapause induction. For example, the RACK (receptor for activated protein kinase) gene appears to be up-regulated in response to diapause-inducing short daylength in *Cabbage armyworm* (Uryu *et al.* 2003). High expression of PP2A-Aα (a structural subunit of the protein phosphatase 2A complex) induced the cotton bollworm, *Helicoverpa armigera* enter facultative pupal diapause during the photoperiod-sensitive stage (Ke and Xu 2013). Transcriptional evidence for sRNA regulation of pupal diapause of the flesh fly, *Sarcophaga bullata*, indicated a role for sRNA in programming the switch from direct development to diapause (Reynolds *et al.* 2013). A global pattern of gene expression associated with very early stages of diapause indicated that short day triggering of diapause was associated with inhibition of 20-HE (20E) signaling during the photoperiod-sensitive period of larvae of the drosophilid fly *Chymomyza costata* (Poupardin *et al.* 2015). Whole-transcriptome microarrays revealed some potential regulatory mechanisms driving diapause induction of *Culex pipiens* female adults, including the TGF-b and Wnt signaling pathways, ecdysone synthesis, chromatin modification, and the circadian rhythm (Hickner *et al.* 2015). In nonblood-fed female adults of *Aedes albopictus*, potential regulatory elements of diapause induction include two canonical circadian clock genes, timeless and cryptochrome1, while in blood-fed females, genes related to energy production and offspring provisioning were differentially expressed, including oxidative phosphorylation pathway and lipid metabolism genes (Huang *et al.* 2015). Global transcriptome analysis provides insight into the foundamental role of the circadian clock in summer diapause induction in onion maggot, *Delia antiqua* (Ren *at al.* 2018). In obligatory diapause insects, a few univoltine insects enter obligatory diapause at specific stages in each generation regardless of the environmental cues it receives. However, little is known about how a diapause induction is regulated in obligatory diapause insects. Therefore, understanding the diapause-inducing mechanism of obligatory diapause insects may enrich the research status of insect diapause and contribute to the in-depth elucidation of the molecular regulation mechanism of insect diapause induction.

Recently, Next-generation sequencing has widely been used to characterize genomes and transcriptomes, especially for insects without reference genome sequences (Ragland *et al.* 2010; Ekblom and Galindo 2011; Liu *et al.* 2014). And next generation sequencing has already led to exciting progress on the transcriptome in several insect species, such as *Bombyx mori* (Xia *et al.* 2004), *Danaus plexippus* (Zhan *et al.* 2011), *Heliconius melpomene* (Consortium 2012) and *Plutella xylostella* (You *et al.* 2013), *Bemisia tabaci* (Wang *et al.* 2010), *Liposcelis entomophila* (Wei *et al.* 2013), *Bactrocera dorsalis* (Shen *et al.* 2011), *Monochamus alternatus* (Lin *et al.* 2015), *Blattella germanica* (Zhou *et al.* 2014), and *Chrysomya megacephala* (Zhang *et al.* 2013), which have been identified some interesting genes and revealed expression patterns and gene function. Three B. minax transcriptome that were previously assembled and annotated can provide several foundations for further DEG analysis (Dong *et al.* 2014a; Wang *et al.* 2016; Wang *et al.* 2017). However, there is no report on diapause induction.

In this study, we used transcriptome sequencing to compare the gene expression profiles of the Chinese citrus furit fly, *B. minax* at second-instar larvae, third-instar larvae and pupal stages, and identified differentially expressed unigenes following diapause using illumina sequencing technology. The results may provide information about potential regulation components of diapause induction for further genomic studies in obligatory diapause insects.

## MATERIALS AND METHODS

### Insect rearing and sampling

Oranges infested with larvae were brought back to the laboratory from an orchard (E 111°42’, N 30°14’) in Songzi County, Jingzhou City, Hubei Province, China, on Oct. 9, 2017. Second-instar larvae (mouth hooks’s length: 0.42-0.61 mm) and third-instar larvae (mouth hooks’s length: 0.65-0.78 mm) collected from the oranges. Some third-instar larvae were placed over sand in plastic dishes and allowed to pupate. All dishes were placed outdoors under natural temperature and light/dark cycle in the Jingzhou district, Jingzhou City, Hubei Province, China. The sand was changed weekly and regularly watered to maintain moisture.

Samples were collected at three stages, second-instar larvae (2L), third-instar larvae (3L) and pupal (P, one week after pupating). Three biological replicates were generated for each stage. All samples were snap frozen in liquid nitrogen and stored at −80°C for subsequent transcriptomic analysis.

### RNA isolation, library construction, and illumina sequencing

Total RNA from each sample was isolated using TRIZOL Reagent (Life technologies, CA, USA) according to the manufacturer’s instructions. RNA degradation and contamination was monitored on 1% agarose gels. RNA concentration and integrity were determined using Qubit^®^ RNA Assay Kit in Qubit^®^2.0 Flurometer (Life Technologies, CA, USA) and the RNA Nano 6000 Assay Kit of the Agilent Bioanalyzer 2100 system (Agilent Technologies, CA, USA). The isolated RNA pellets were stored at −80°C until needed. Sequencing libraries were generated using NEBNext^®^ Ultra™ RNA Library Prep Kit for Illumina^®^ (NEB, USA) following manufacturer’s recommendations and index codes were added to attribute sequences to each sample. The clustering of the index-coded samples was performed on a cBot Cluster Generation System using TruSeq PE Cluster Kit v3-cBot-HS (Illumia) according to the manufacturer’s instructions. After cluster generation, the library preparations were sequenced on an Illumina Hiseq 2000 platform and paired-end reads were generated.

### Raw data colledtion, assembly, and annotation

The raw reads of fastq format were firstly processed through in-house perl scripts, and clean reads were obtained by removing reads containing adapter, reads containing ploy-N and low quality reads from raw data. All the downstream analyses were based on clean data with high quality. Transcriptome assembly was accomplished based on the left.fq and right.fq using Trinity (Grabherr *et al.* 2011) with min_kmer_cov set to 2 by default and all other parameters set default. Assembled unigenes were used for annotating based on the following database: NR (NCBI non-redundant protein sequences); GO (Gene Ontology); COG (Clusters of Orthologous Groups of proteins); KEGG (Kyoto Encyclopedia of Genes and Genomes).

### DEGs analysis

Gene expression levels were estimated by RSEM (Li *et al.* 2011) for each sample. Clean data were mapped back onto the assembled transcriptome and read count for each gene was obtained from the mapping results. Differential expression analysis of two groups was performed using the DESeq R package (1.10.1). DESeq provide statistical routines for determining differential expression in digital gene expression data using a model based on the negative binomial distribution. The resulting P values were adjusted using the Benjamini and Hochberg’s approach for controlling the false discovery rate (FDR). Genes with an adjusted FDR <0.001 and |log_2_ FC (fold change)|≥2 were assigned as differentially expressed.

### Function and pathway enrichment analysis of DEGs

For pathway enrichment analysis, all of the differentially expressed genes were mapped to GO and KEGG pathway terms and the significantly enriched terms were filtered. GO enrichment analysis of DEGs was implemented by the topGO R packages based Kolmogorov-Smirnov test (Ashburner 2000), and used KOBAS (Xie *et al.* 2005) software to test the statistical enrichment of differential expression genes in KEGG pathways.

### Validation of RNA-Seq result by qRT-PCR

qRT-PCR was performed to verify the accuracy of the differentially expressed genes analysis. RNA sample from the three developmental stages were the same as those used for RNA-Seq. Total RNA was reverse transcribed into cDNA using SYBR^®^ Premix DimerEraser™ (perfect Real Time) Kit (Takara, Shiga, Japan). Six pairs of specific primers were designed to amplify the genes selected from multiple comparisons (Table S1). *Ubiquitin* was used as a reference gene for normalization (Wang *et al.* 2014). qRT-PCR was conducted in 25μL volumes containing 12.5μL SYBR Premix DimerEraser (2x) 2μL primers (10μM), 1μL cDNA, and 9.5μL ddH_2_O, using a CFX96^TM^ Real-Time PCR Detection System thermal cycler (BIO-RAD, Hercules, CA, USA). Amplification conditions were as follows: initial denaturation at 95°C for 30s; followed by 40 cycles of denaturation at 95°C for 5s, 60°C for 30s. Pearson’s r correlation coefficient was calculated to evaluate the correlation between the qRT-PCR and DEG data. Three biological and three technical replicates were performed for each gene.

### Data availability

The raw data produced in this study have been deposited at NCBI systerm under project number PRJNA545883. BioSample number 2th instar larva-1: SAMN12011777; 2th instar larva-2: SAMN12011778; 2th instar larva-3: SAMN12011779; 3th instar larva-1: SAMN12011780; 3th instar larva-2: SAMN12011781; 3th instar larva-3: SAMN12011782; pupal-1: SAMN12011783; pupal-2: SAMN12011784; pupal-3: SAMN12011785. Other data are within the paper and its supplemental files.

## RESULTS

### Illumina sequencing and data processing

Nine mRNA libraries, three biological replicates for each developmental stage, were sequenced. A total of 72.04G raw reads were generated in all libraries. After removing low quality sequences and ambiguous nucleotides, 240,721,613 clean reads were obtained (Table 1). The number of clean reads ranged from 24,190,185 to 30,582,464, and the ratio of mapped reads exceed 80.64% in all libraries (Table S2). The transcripts were further assembled into 116,402 unigenes with a mean length of 858.16bp (Table 1). Of these unigenes, 91,069 (78.24%) were 200-1000bp in length and 10,474 (9.00%) were > 2000bp, with most unigenes falling between 200bp and 500bp (55.66%) (Figure 1).

**Figure 1.**
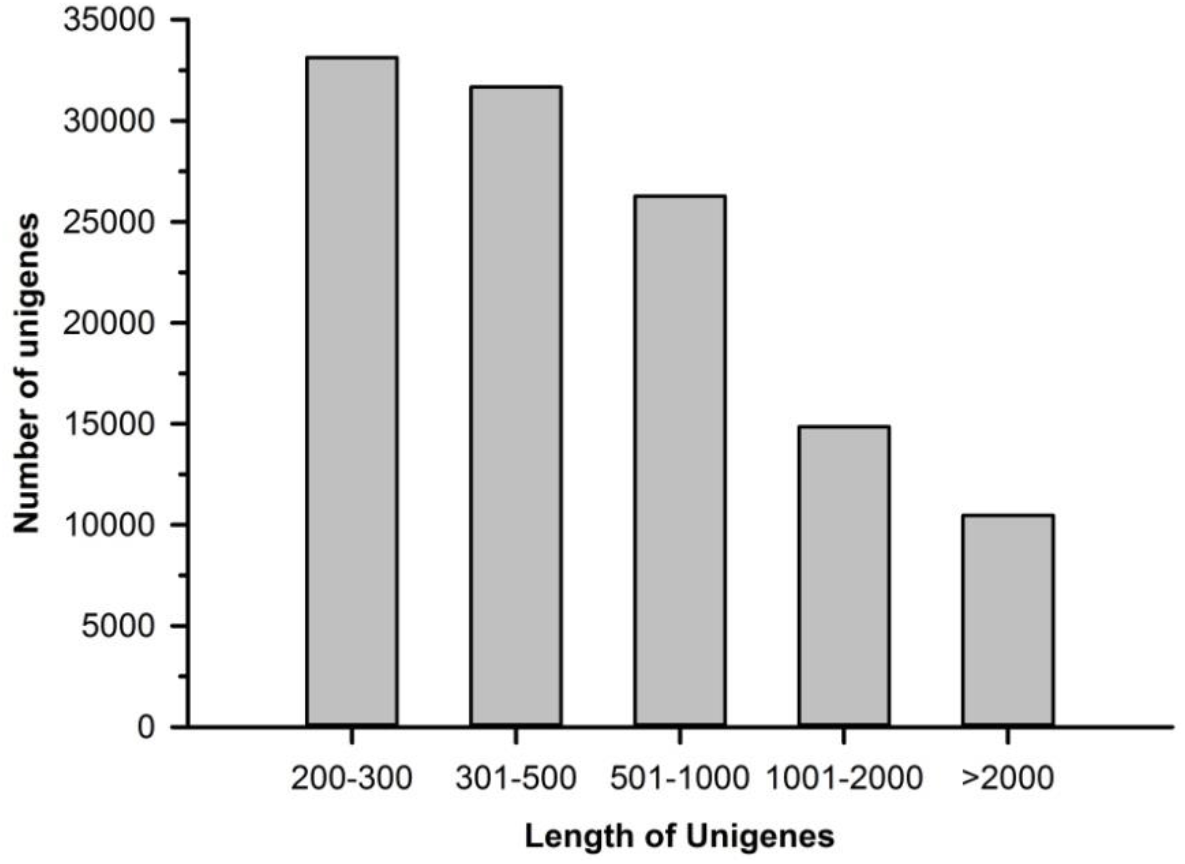
Length distribution of unigenes. A total of 116,402 unigenes were assembled.

**Table 1.** Summary of RNA sequencing, assembling, and functional annotation for *B. minax*

| Sequencing and Assembling | Statistion and Annotation (E-value $\leq 1e-5$) |
| --- | --- |
| Raw Reads (G) | 72.04 |
| Clean Reads Number | 240,721,613 |
| Total nucleotides (nt) | 72,041,000,718 |
| GC Percentage of Total Clean Nucleotides | 42.01% |
| Number of Unigenes | 116,402 |
| Total Length (nt) of Total Unigenes | 99,892,023 |
| Mean Length (nt) of Total Unigenes | 858.16 |
| N50 (nt) of Unigenes | 1472 |
| Unigenes with Nr Database | 44,274 (38.04%) |
| Unigenes with Swiss-Prot Database | 23,724 (20.38%) |
| Unigenes with KEGG Database | 22,366 (19.21%), 295 pathways |
| Unigenes with COG Database | 18,833 (16.18%), 24 functional categories |
| Unigenes with GO Database | 21,966 (18.87%) |
| Biological Process | 20 subcategories |
| Cellular Component | 19 subcategories |
| Molecular Function | 19 subcategories |
| Total Unigenes Annotated | 54,781 (47.06% of 116,402) |

### Annotation of unigenes

Of all unigenes, 54,781 (47.06%) unigenes were successfully annotated (Table 1). A total of 44,274 (38.04%) unigenes were annotated in Nr database, because the genome sequence of *B. minax* has not been reported, sequence alignment of the experimental unigenes was performed using the known genomes of other species. In the species distribution showed that genes from *B. minax* had the greatest number of matches with those of the *Bactrocera dorsalis* (5,837, 13.2%), followed by *Bactrocera cucurbitae* (4,968, 11.23%) (Figure 2).

**Figure 2.**
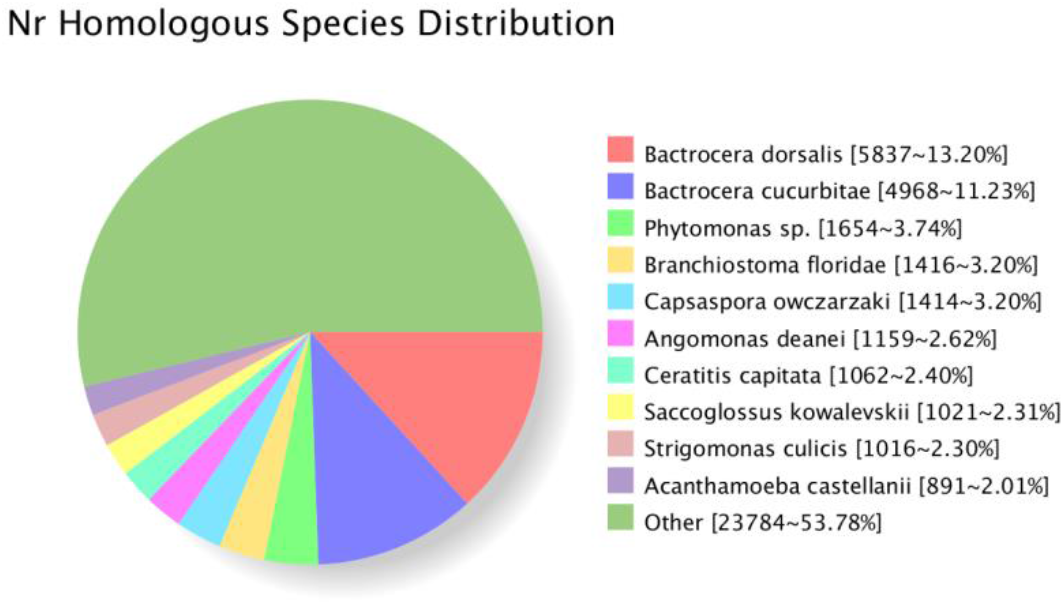
Homology search against NR database for *B. minax* transcriptome unigenes.

GO is a standardized gene functional classification system that provides a structured and controlled vocabulary to predict gene function (Ashburner *et al.* 2000). In this experiment, 21,966 (18.87%) unigenes were grouped into 58 GO functional categories, which were distributed under three categories of Biological Process (n=20), Cellular Components (n=19) and Molecular Function (n=19) (Figure 3). Among biological process, metabolic process, single-organism process and cellular process were the top 3 abundant groups. In term of molecular function, the catalytic activity category was the most abundant, followed by the binding and transporter activity categories. Among the cellular components, the cell, cell part and organelle accounted for the majority of unigenes in unigene classification (Figure 3).

**Figure 3.**
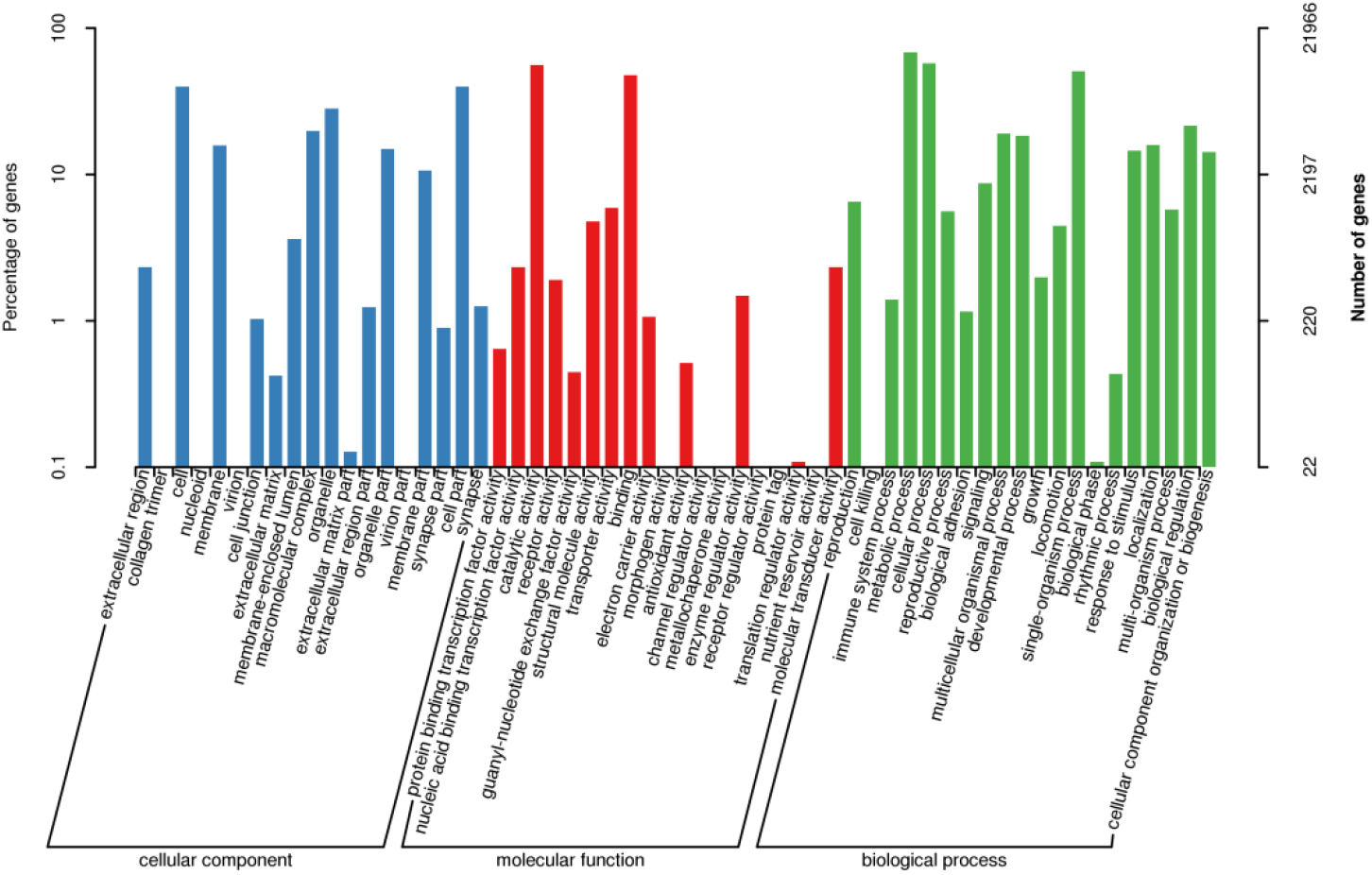
GO classification of *B. minax* transcriptome unigenes.

To analyze the integrity of the libraries and the effectiveness of the annotation process, COG functional classification was performed on the unigene alignment with the COG database using gapped blast and PSI blast program (Altschul 1997). A total 18,833 unigenes were annotated to 24 COG categories (Figure 4). The largest group in the cluster was “general function prediction only”, with 4848 unigenes; followed by “translation, ribosomal structure and biogenesis”, with 2533 unigenes and “amino acid transport and metabolism”, with 2256 unigenes.

**Figure 4.**
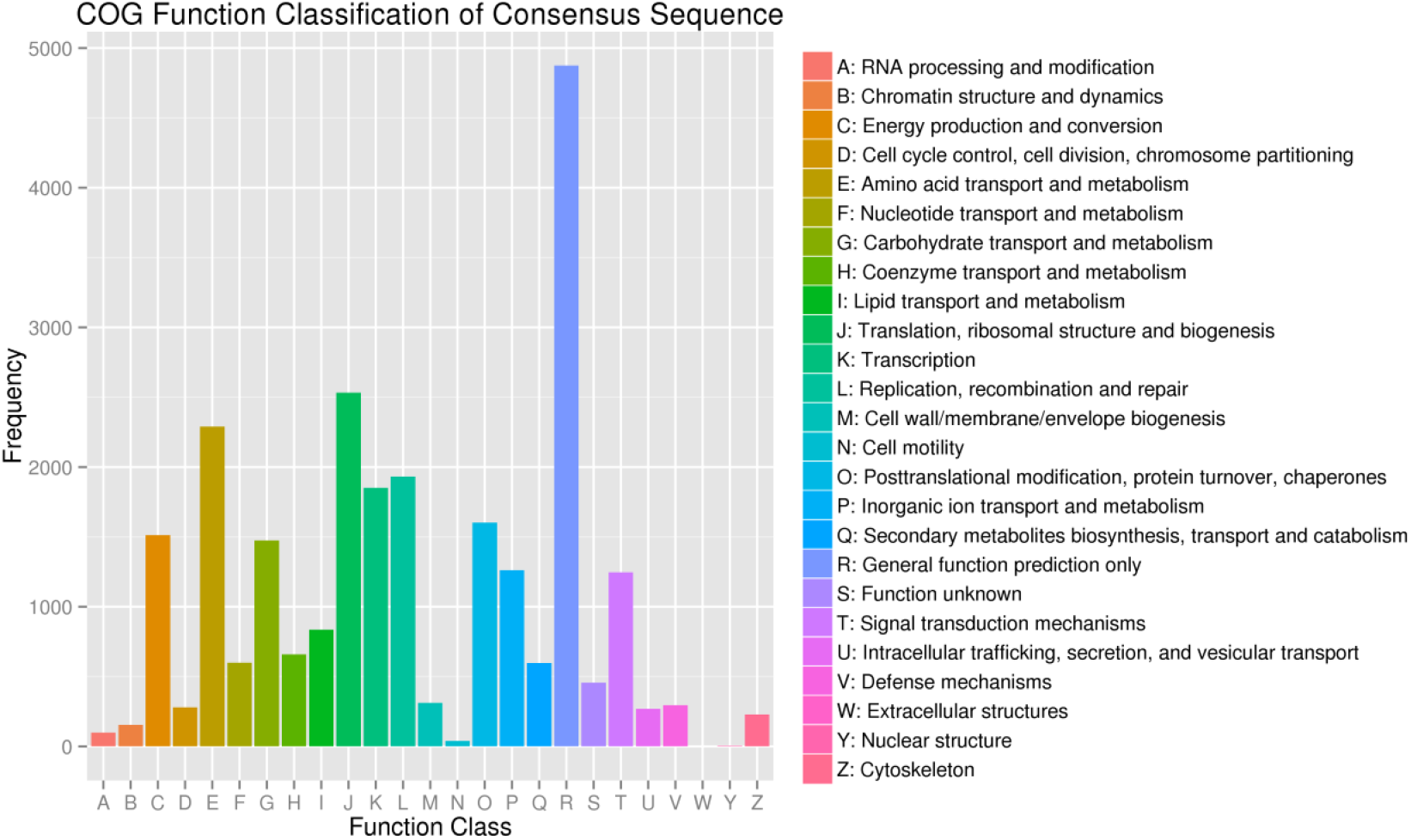
COG classification of *B. minax* transcriptome unigenes.

The KEGG pathway assignment was also performed for all assembled unigenes to categorize gene functions, focusing on biochemical pathways (Kanehisa and Goto, 2000). A total of 22,366 unigenes were annotated against the KEGG database and were assigned to 295 pathways (Table 1). Among these pathways, ribosome, carbon metabolism and protein processing in endoplasmic reticulum were the most represented, with 1002 unigenes, 878 unigenes and 669 unigenes, respectively (Table S3). We identified the areas of interest to further analyze these annotations, providing a valuable resource for elucidating functional genes in pupal diapause induction of *B. minax*.

### Analysis of gene expression profies

To identify significant expression changes in genes, we conducted a differential expression analysis of unigenes expression through pairwise comparisons of the second-instar larvae (2L), third-instar larvae (3L) and pupal (P, one week after pupating). A total of 9,934 unigenes were significantly differentally expressed in three pairwise comparisons (Figure S1). All differentally expressed genes were divided into 6 groups with each exhibiting a representative expression pattern. Genes in group C and D were highly expressed in pupal stage, whereas genes in other groups were lowly expressed in pupal stage (Figure 5). These results shown that most unigenes were silent may due to the slow pace at which physiological activities and growth occur when larva entering pupal stage and entering pupal obligatory diapause.

**Figure 5.**
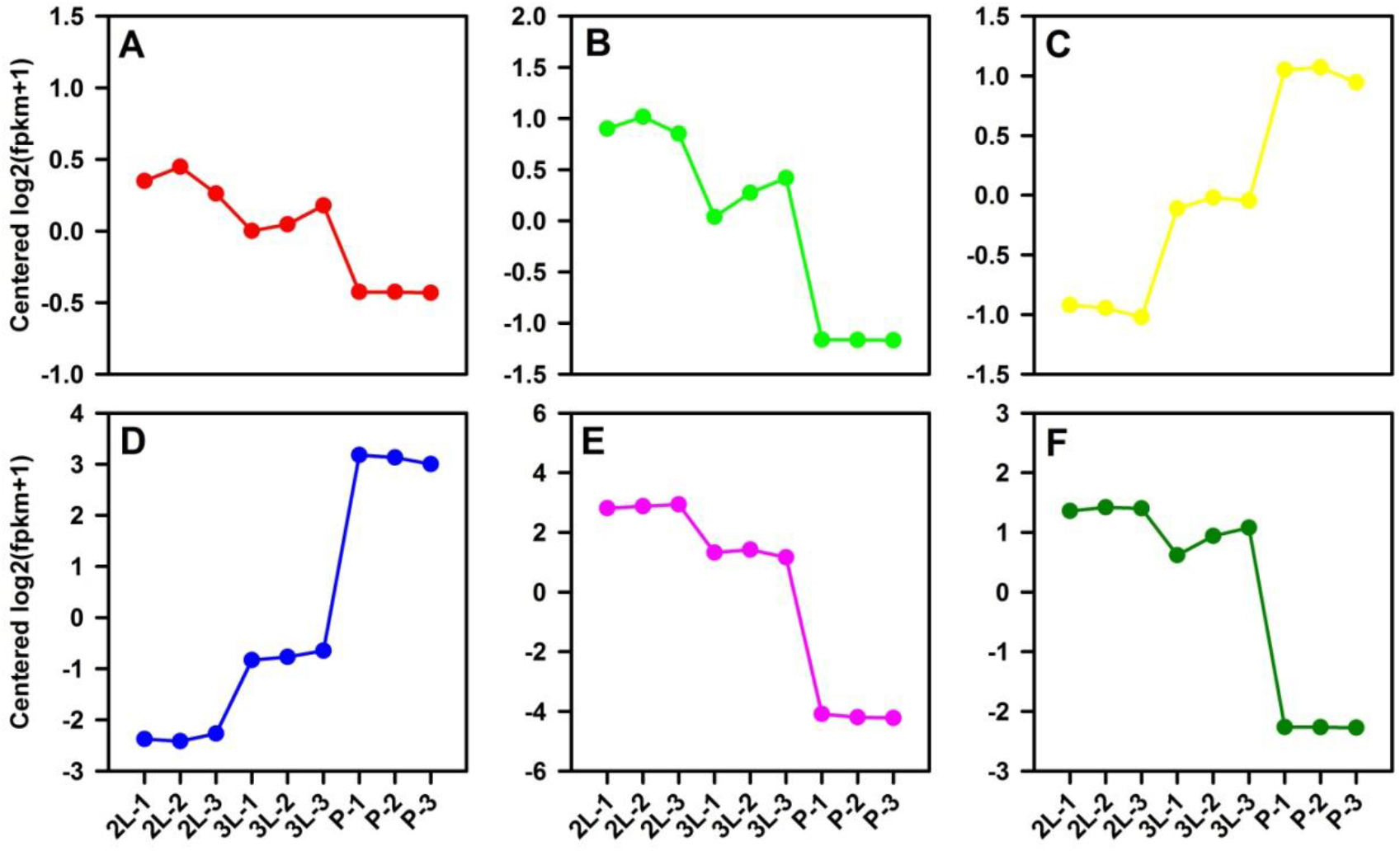
Groups of differentially expressed genes (DEGs) among three different *B. minax* developmental stage.

### Functional enrichment analysis for DEGs

To understand the functions of the differentially expressed genes, we compared the GO term associated with the three different stages after mapping all the DEGs to the GO database. According to the GO classification, most unigenes were associated with metabolic process, catalytic activity, cell, cell part, single-organism process, binding and cellular process (Figure 6), the metabolic process was the most highly represented category, which led to in-depth analysis of this group. The top 5 significantly enriched term for each compares were list in Table S4.

**Figure 6.**
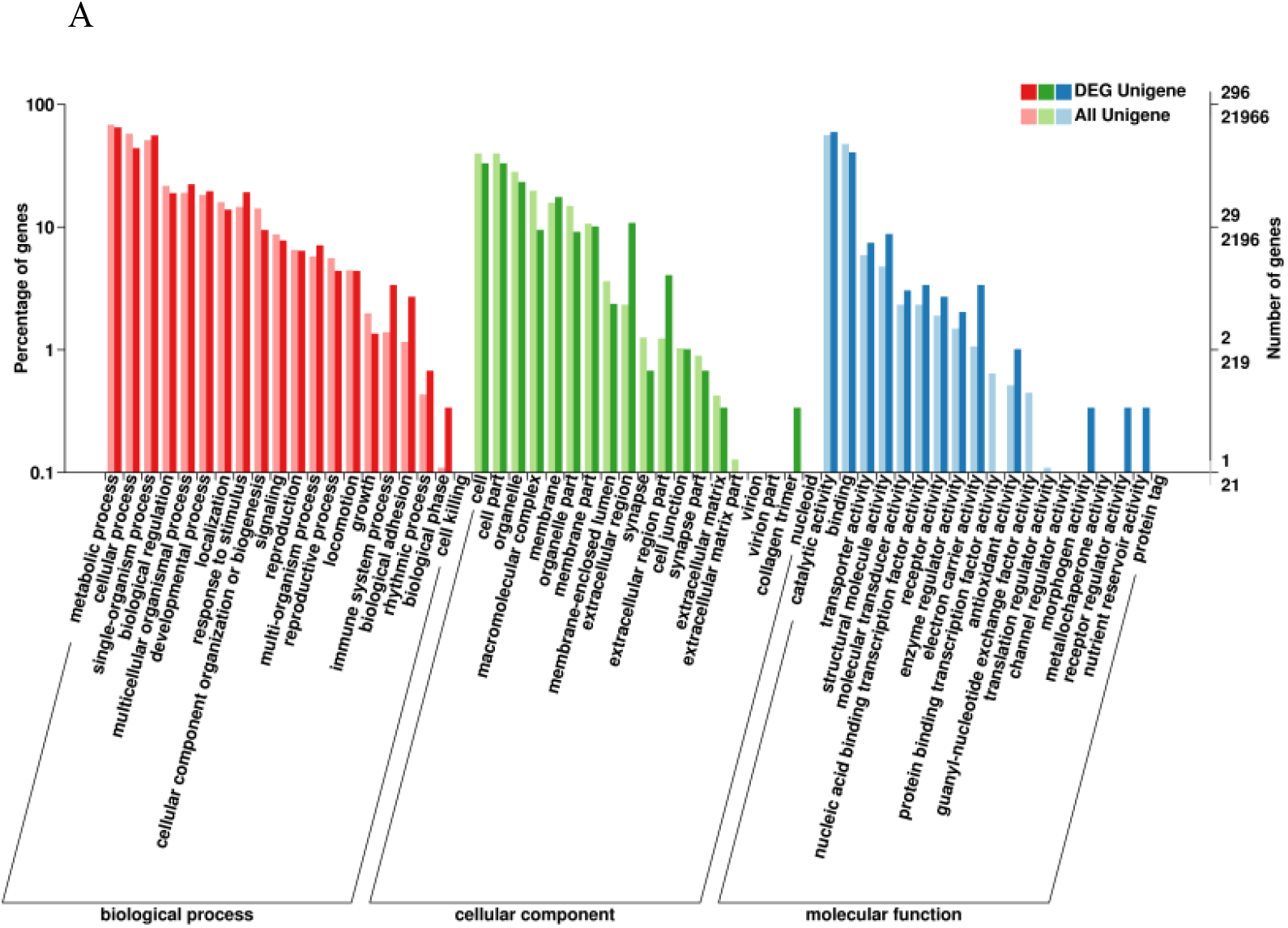
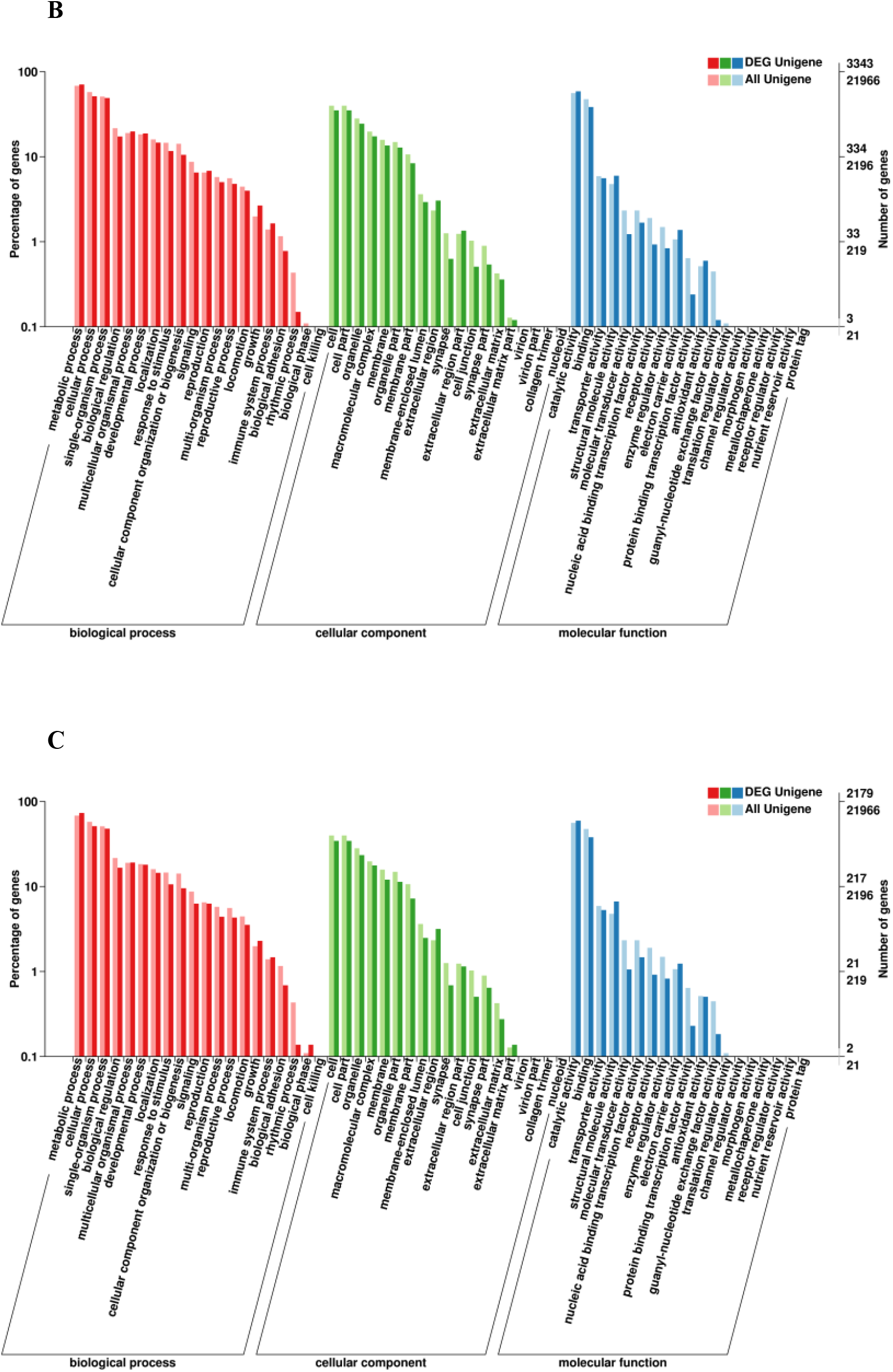
GO annotation of differentially expressed genes in 3L vs. 2L (A), P vs. 2L (B), P vs. 3L (C). Left panel, the y-axis indicate the percentage of a specific category of unigenes; right panel, the y-axis indicates the number of unigenes in a category.

KEGG pathway enrichment analysis showed that 41 pathways were significantly enriched with corrected *P* value ≤ 0.05 in 3L vs 2L, P vs 2L and P vs 3L. All of the significant pathways are listed in Table S5. Of these, in 3L vs 2L, most DEGs were classified into pyruvate metabolism, glycolysis / gluconeogenesis and biosynthesis of amino acids. In P vs 2L, most DEGs were assigned to biosynthesis of amino acids, carbon metabolism and glycolysis / gluconeogenesis. And most DEGs were classified into ribosome, biosynthesis of amino acids and carbon metabolism in P vs 3L. These results shown that, in the developmental process from larval to pupal, most DEGs were related to biosynthesis of amino acids, carbon metabolism and glycolysis / gluconeogenesis. This may release that those DEGs were related to diapause induction in the Chinese citrus fruit fly.

### Validation of RNAseq results using qRT-PCR

To validate the RNAseq results by illumina sequencing, the 6 DEGs in three different compares were validated throught quantitative real-time PCR. The results showed a strong correlation between the qRT-PCR and DEG date with Pearson’s correlation coefficient > 0.99 (Figure 7), indicating the reliability of using DEG date to investigate temporal-specific gene expression profiles at the three stages.

**Figure 7.**
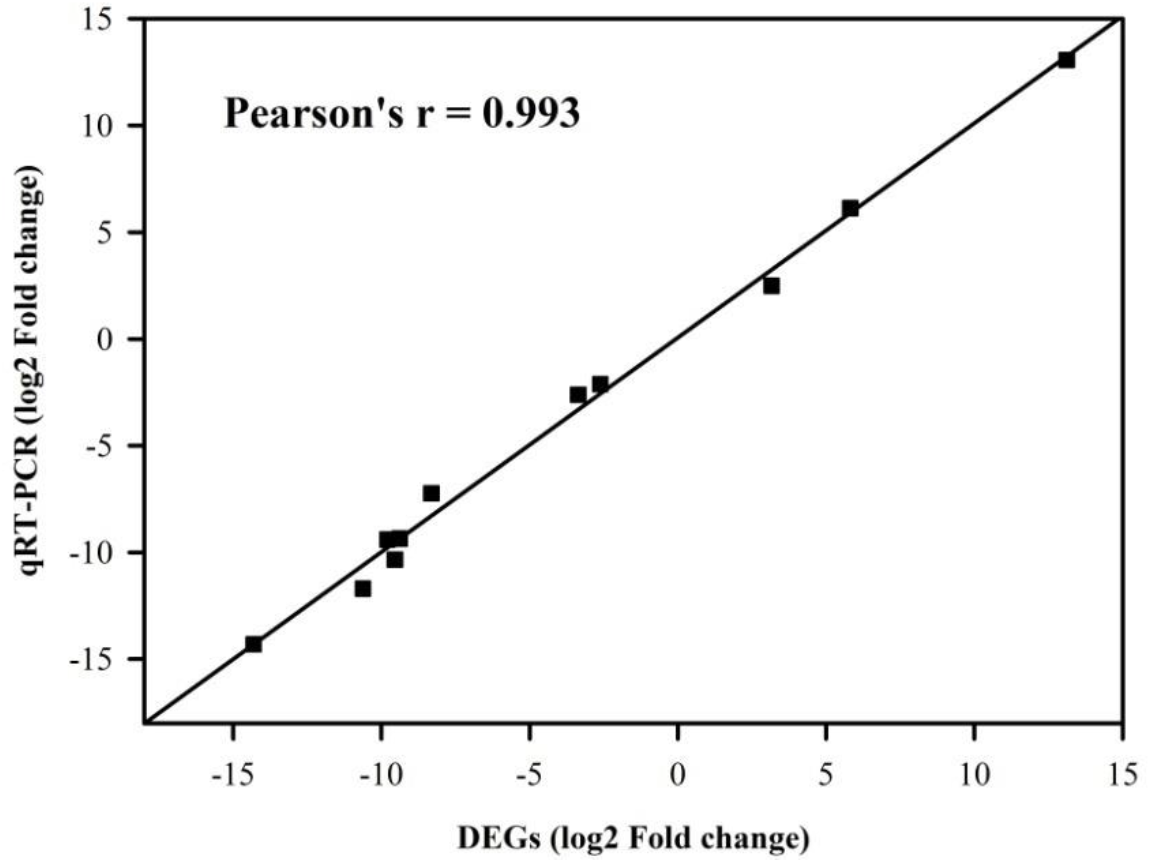
Correlation analysis of qRT-PCR and differentially expressed gene (DEG) date for selected genes of *Bactrocera minax.*

## DISCUSSION

Obligatory diapause is not elicited by environmental cues because it represents a fixed component of ontogenetic programme and is expressed regardless of the environmental condition (Koštál 2006). In the development process, obligate diapause insects enter into the diapause state when they enter a specific stage (Koštál, 2006). Therefore, diapause insects enter into obligatory diapause may be the result of specific expression of particular gene at specific time. Under the natural environment, after larval pupation, *B. minax* enters into obligatory diapause at pupal stage. From the previous one to the diapausing stage, significant differential expression genes may be the potential regulators of inducing obligatory diapause. According to the comparing of the transcriptome between diapause and non-diapause, all DEGs were divided into 6 groups (Figure S2). Throughout 2L-3L-P developmental axis, the expression of genes in group B, E and F were suppressed, whereas those genes in group C and D were activated. Therefore, those genes were highly or lowly expressed in pupal stage may relate to obligatory diapause induction in the Chinese citrus fruit fly, *Bactrocera minax*.

It is well known that the endocrine hormones control the diapause program (Denlinger *et al.* 2011). The prothoracicotropic hormone (PTTH) receptor signaling transduction (Young *et al.* 2012), Juvenile hormone and ecdysone biosynthesis (Denlinger *et al.* 2011) are closely related to diapause, which involves several KEGG pathways, including MAPK signaling pathway (Ko04010), Wnt signaling pathway (Ko04310), mTOR signaling pathway (Ko04150), Calcium signaling pathway (Ko04020), Steroid biosynthesis (Ko00100), Steroid hormone biosynthesis (Ko00140), Terpenoid backbone biosynthesis (Ko00900), Insect hormone biosynthesis (Ko00981), FoxO signaling pathway (Ko04068) and Insulin signaling pathway (Ko04910). Many unigenes belonging to these pathways were identified in *B. minax* transcriptome. The KEGG pathway assignment will be helpful for predicting the functions of *B. minax* genes, and will contribute to the further research on relevant diapause initiation and termination.

In all arthropods, the ecdysteroids mediate transitions between developmental stages (Gilbert *et al.* 2003). The ecdysteroids are also very central in regulating many forms of insect diapause (Denlinger 2002; Denlinger *et al.* 2012; Denlinger 2000). The prohormone ecdysone is synthesized from dietary cholesterol or phytosterols. In larval stages of insects, the biosynthetic pathway is localized in the prothoracic gland (part of a ring gland in larval drosophilids). Ecdysone is released by ring gland and further converted into the active hormone 20-hydroxyecdysone (20E) in target tissues (Gilbert *et al.* 2002; Yamanaka *et al.* 2013). The changes of ecdysteroid titer have been recognized from 3rd larval instar to pupal of *B. minax* (Wang *et al.* 2014). After larval pupation, ecdysteroid titer decreased significantly. During pupal stage, ecdysteroid titer increase as the time of pupal developmental. Additionally, 20E could break pupal diapause of *B. minax* by topical application (Chen *et al.* 2016; Wang *et al.* 2014). Therefore, we speculated that the low level of ecdysteroid titer inhibited the developmental of the pupal, which led to obligatory diapause in pupal stage of *B. minax.* Moreover, ecdysteroid regulated the induction and maintenance of the pharate first instar diapause of the gypsy moth, Lymantria dispar, which is obligatory diapause (Lee and Denlinger 1997; Lee *et al.* 1997). Therefore, the synthesis and release of ecdysteroid may regulate potentially the induction of obligatory diapause of *B. minax.*

Most of ecdysteroid biosynthetic enzymes belong to the family of cytochromes P450 (Niwa 2010; Pondeville 2013). Halloween genes are a set of genes encoding cytochrome P450 enzymes, including Spook (CYP307a1), Spookier (CYP307a2), Phantom (Cyp306a1), Disembodied (Cyp302a1), Shadow (Cyp315a1), and Shade (Cyp314a1) (Kankare *et al.* 2010; Petryk *et al.* 2003; Warren *et al.* 2004; Yoshiyama *et al.* 2006). The expression pattern of those Halloween genes was list in Table S6. Only Cyp314a1 was significantly down-regulated in P vs 3L. This results suggest that inhibition of the 20E biosynthetic pathway (downregulation of Shade/Cyp314a1 expression), might represent important early steps in diapause induction in *B. minax* pupal. Our speculation was indirectly supported by the transcriptomic date. Important signaling pathway

By definition, insect diapause is a centrally regulated arrest, or significant slowdown, of development (Denlinger 2002; Koštál 2006). In pupae of *B. minax*, the arrest of development is obviously expressed as a significant slowdown/cessation of the tissue differentiation (Chen *et al.* 2016). Previous research has shown that the arrest of cell cycle (Ko04110) is a hallmark of diapause in insects (Koštál *et al.* 2009). Based on KEGG enrichment analysis, there were 19 DEGs of cell cycle, all of which are down-regulation (Table S7). Our results suggest that down-regulation of nine DEGs related to cell cycle control, which makes it a good candidate for mediation of the inhibition of cell cycle in response to diapause induction of obligatory diapause. The MCM (minichromosome maintenance) family of proteins contributes to the initiation and competent state of DNA replication (Pucci *et al.* 2007). According to our results, down-regulation of two DEGs (c57934. graph_c0 and c59738.graph_c0) encoding MCM in cell cycle may inhibit DNA replication in the Chinese citrus fruit fly, and result in cell cycle arrest during diapause induction.

Juvenile hormones (JHs) are acyclic sesquiterpenoids that regulate many aspects of insect physiology, including development, reproduction, and polyphenisms (Riddiford 1994; Wyatt and Davey 1996), and play key roles in insect diapause (Rinehart *et al.* 2007; Salvucci *et al.* 2000; Yagi 1976). The insulin can regulate the synthesis of juvenile hormone of *Culex pipiens* to mediate the diapause response, and in the diapause period, the expression of insulin signaling leads to the down-regulation of JH and up-regulation of fork head transcription actor (FOXO) to promote fat hypertrophy (Sim and Denlinger 2008). In some insect species, insulin signaling pathway even involves regulation of the diapause phenotype (Sim and Denlinger 2013). In the Chinese citrus fruit fly, we found one DEG in 3L VS 2L, twenty eight DEGs in P VS 2L and twenty DEGs in P VS 3L, and those DEGs ralated to insulin signaling pathway. And also, in FoxO signaling pathway, five DEGs in 3L VS 2L, fifty eight DEGs in P VS 2L and thirty two DEGs in P VS 3L. We speculate that those DEGs in this two pathway maybe arrested in obligatory diapause induction of *B. minax*, contributing to induce diapause.

GO and KEGG enrichment analysis indicates that most of the up-regulated and down-regulated genes are involved in metabolic process (biological process) and metabolic pathway (Table S4 and Table S5). Cross talk between the brain and fat body as a regulator of diapause suggested that the TCA cycle may be a checkpoint for regulating insect diapause (Xu *et al.* 2012). 45 DEGs related to TCA cycle are involved in energy production and conversion, amino acid transport and metabolism and carbohydrate transport and metabolism, were differentially down-regulated during larval-pupal period (Table S8). These patterns indicated a metabolic switch during diapause induction, and some candidate genes were revealed may as potential regulators of obligatory diapause in *B. minax.*

Our study is the first evaluation of the molecular mechanisms of obligatory diapause induction in the Chinese citrus fruit fly, *B, minax*. We report compelling differences between diapause and non-diapause (before diapause) populations that will enhance our understanding of the molecular of mechanisms of obligatory diapause induction, and further our understanding of the biology and ecology of the Chinese citrus fruit fly.

## ACKNOWLEDGMENTS

We thank Dr. Junliang Yin for his assistance in uploading raw data of transcriptome to NCBI system. This research was supported by the National Natural Science Foundation of China (31572010). The authors declare no conflicts of interest.

